# EPP1 couples receptor activation to cytoplasmic signaling in root nodule symbiosis

**DOI:** 10.1101/2025.09.29.679184

**Authors:** Malita M. M. Nørgaard, Nikolaj B. Abel, Emma Birkeskov, David Landry, Jens Stougaard, Kasper Røjkjær Andersen

**Author notes:** These authors contributed equally to this work (Malita M. M. Nørgaard, Nikolaj B. Abel).

## Abstract

Legume symbiosis with nitrogen-fixing bacteria is controlled by a cascade of signaling events leading to root nodule development. While plant cell-surface receptors initiate this process, the link between receptors and cytoplasmic signaling components remains unclear. Here we identify Early Phosphorylated Protein 1 (EPP1) as a central mediator of this pathway. EPP1 is recruited to the activated SYMRK receptor, where it is phosphorylated on a key serine residue, an event essential and sufficient to propagate symbiotic signaling. We provide structural and functional validation of the SYMRK-EPP1 signaling complex and demonstrate that EPP1 is required for root nodule formation. Synthetic engineering of the SYMRK-EPP1 interaction bypasses the need for symbiotic bacteria to initiate the pathway and triggers nodule organogenesis. These findings establish EPP1 as a crucial cytoplasmic component between receptor activation and intracellular signaling, advancing our understanding of nitrogen-fixing symbiosis.

## Introduction

Legumes form symbiotic relationships with nitrogen-fixing bacteria, enabling plant growth in nitrogen-poor environments^1^. In *Lotus japonicus* (*Lotus*), this interaction is initiated by recognition of bacterial lipochitooligosaccharide signals, known as Nod factors, by Nod factor receptors NFR1 and NFR5^2–7^. The NFRs relay the symbiotic signal to the malectin-like/leucine-rich repeat receptor kinase SYMRK^8,9^, which is essential for downstream signaling in root nodule symbiosis. Recent findings show that phosphorylation of four conserved residues on the SYMRK alpha-I motif is critical for its activation and function^10^. Once activated, SYMRK triggers a signaling cascade leading to oscillatory calcium spiking in the nucleus. These calcium signals are decoded by the calcium- and calmodulin-dependent protein kinase CCaMK^11,12^, which activates the transcription factor NODULE INCEPTION (NIN) to drive nodule organogenesis and bacterial infection^13,14^. While this pathway has been mapped at the receptor- and transcriptional levels, the cytoplasmic components that couple receptor activation to nuclear calcium spiking remain poorly understood. Early Phosphorylated Protein 1 (EPP1) was identified as a Nod factor-responsive phospho-protein^15^ and subsequent studies showed that EPP1 regulates the expression of symbiotic genes like NIN, ERN1 and NF-YB^16,17^. However, its function in symbiosis has remained unresolved. In this study, we demonstrate that EPP1 is an essential signaling component in root nodule symbiosis in *Lotus*. We show that EPP1 is directly recruited to SYMRK in a phosphorylation-dependent manner, acting downstream of SYMRK activation. Using structural modeling, *in vitro* binding, phosphorylation assays, and genetics, we reveal that SYMRK phosphorylates EPP1 on a conserved serine residue and that this modification is required for nodule formation. Strikingly, we find that synthetic recruitment of EPP1 to SYMRK is sufficient to trigger spontaneous nodulation in the absence of rhizobia, positioning EPP1 as a critical signaling hub linking SYMRK activation to the initiation of intracellular symbiotic signaling.

## Results

### EPP1 is essential for root nodule symbiosis

In *Lotus*, EPP1 is encoded by two paralogous genes, *Epp1a* and *Epp1b*. We obtained two independent single-allele knockout LORE1 lines for each gene: two for *epp1a* and two for *epp1b* and crossed these to obtain two independent *epp1ab* double mutant lines. All single *epp1* mutant plants were symbiotically competent and formed infected red nodules when inoculated with *Mesorhizobium loti* (*M. loti*), the symbiont of *Lotus* (Fig. 1a-b). We observed a small reduction in the number of red nodules compared to Gifu wildtype, but all single *epp1* mutant plants formed infection threads similar in both number and morphology to wildtype plants (Fig. 1c-d). However, we observed a clear symbiotic phenotype for the *epp1ab* double mutant plants, as all were unable to form infection threads and nodules (Fig. 1a-d), showing that EPP1 is essential in *Lotus*. To demonstrate that *Epp1* is the causative gene for this observed phenotype, we next performed hairy root complementation in the *epp1ab* double mutant plants. Using the native promoters to drive the expression of either EPP1a or EPP1b restores nodulation (Fig. 1e), revealing a level of redundancy between the two genes. As expression of *Epp1a* is approximately eight times higher than *Epp1b* in root hairs (Extended Data Fig. 1), we decided to focus the study on EPP1a for its role in root nodule symbiosis.

**Figure 1.**
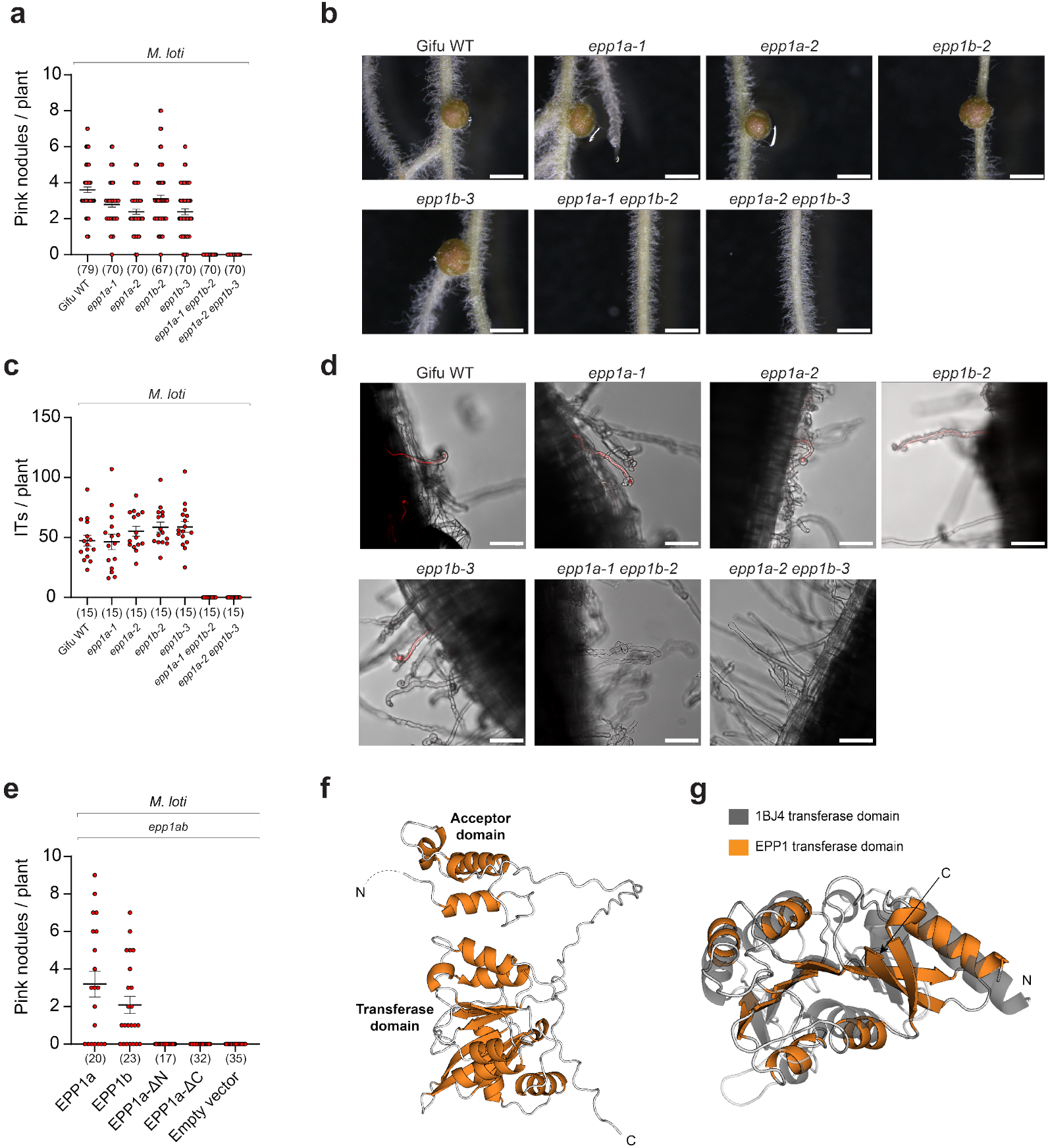
EPP1 is required for nodule organogenesis and rhizobial infection. **a.** Number of pink infected nodules on indicated mutants 4 weeks after inoculation with *M. loti* rhizobia. The numbers below graph specify the number of plants for the specific mutant analyzed. **b**. Representative pictures of nodules from (**a**). Scalebar indicates 1 mm. **c**. Number of infection threads on indicated mutants 10 days after inoculation with *M. loti* rhizobia. The numbers below graph specify the number of plants for the specific mutant analyzed. **d**. Representative pictures of root hairs with infection threads or swollen tips from (**c**). Scalebar indicates 80 µm. **e**. Number of pink infected nodules 4 weeks after inoculation with *M. loti* rhizobia on *epp1a-1 epp1b-2* (*epp1ab*) mutants expressing indicated versions of EPP1 driven by its native promoter. The numbers below graph specify the number of plants for each version of EPP1 or empty vector. **f**. AlphaFold3 model of the predicted domain structure of *Lotus* EPP1a. **g**. The transferase domain of *Lotus* EPP1a (orange) is superimposed on the major catalytic domain of PDB entry 1BJ4 (gray), revealing conserved architecture characteristic of a fold-type I PLP-dependent enzyme (RMSD = 3.5 Å).

To understand the domain structure of the EPP1 protein, we modelled EPP1a using AlphaFold3^18^ (Fig. 1f). EPP1a consists of a disordered N-terminal region (residues 1-58), followed by a helix-turn-helix domain (residues 59-115) hereafter referred to as the ‘acceptor domain’ as it was shown to be phosphorylated in *Medicago truncatula* EPP1 upon Nod factor treatment^15^. The acceptor domain is connected via a flexible linker (residues 116-161) to the C-terminal ‘transferase domain’ (residues 162-364), named after its structural similarity to the fold type I family of pyridoxal 5’-phosphate (PLP)-dependent transferases (Fig. 1g)^19^. To test that both domains are necessary for EPP1 function, we complemented the *epp1ab* double mutant with EPP1 versions that lack one or the other domain and find that both variants fail to restore nodulation (Fig. 1e and Extended Data Fig. 2). Together, our genetic data demonstrate that EPP1 is an essential component in root nodule symbiosis and that both the acceptor and transferase domains have critical functions.

### SYMRK and EPP1 form a stable complex

The kinase activity of SYMRK is essential for signal propagation, but its substrate that propagate signaling is still unknown. To place EPP1 within the symbiotic pathway, we therefore tested whether EPP1 interacts directly with SYMRK. We expressed and purified *Lotus* EPP1a (hereafter EPP1) and two different versions of the *Lotus* SYMRK kinase (Extended Data Fig. 3). We produced a gain-of-function version of SYMRK 4D, which is an active form of SYMRK where four conserved phosphorylation sites in the alpha-I motif (S877, S885, S889 and S893) are phospho-mimicked by aspartates^10^. We pre-incubated EPP1 and the activated form of SYMRK 4D in a 1:1 molar ratio and analyzed complex formation using analytic gel filtration. Individually, EPP1 and SYMRK 4D migrated in single peaks with apparent molecular weights of ∼70 kDa and ∼50 kDa respectively (Fig. 2a). In contrast, pre-incubating EPP1 and SYMRK 4D resulted in the formation of a large stable complex on gel filtration of ∼100 kDa (Fig. 2a). To investigate if the phosphorylations of the alpha-I motif are important for the interaction, we repeated the experiment with a SYMRK 4A version with alanine replacements at these positions. SYMRK 4A and EPP1 fail to form a stable complex on gel filtration as the pre-incubated complex migrates as two distinct peaks that correspond to each individual protein peak (Fig. 2b). We next evaluated complex formation using full length receptors and performed co-immunoprecipitation with SYMRK variants and EPP1 WT in *Nicotiana benthamiana* leaves. This experiment shows that EPP1 is strongly enriched only with SYMRK 4D and not with SYMRK or SYMRK 4A (Extended Data Fig. 4). Together, our data show that only the activated and phosphorylated state of SYMRK forms a stable complex with EPP1.

**Figure 2.**
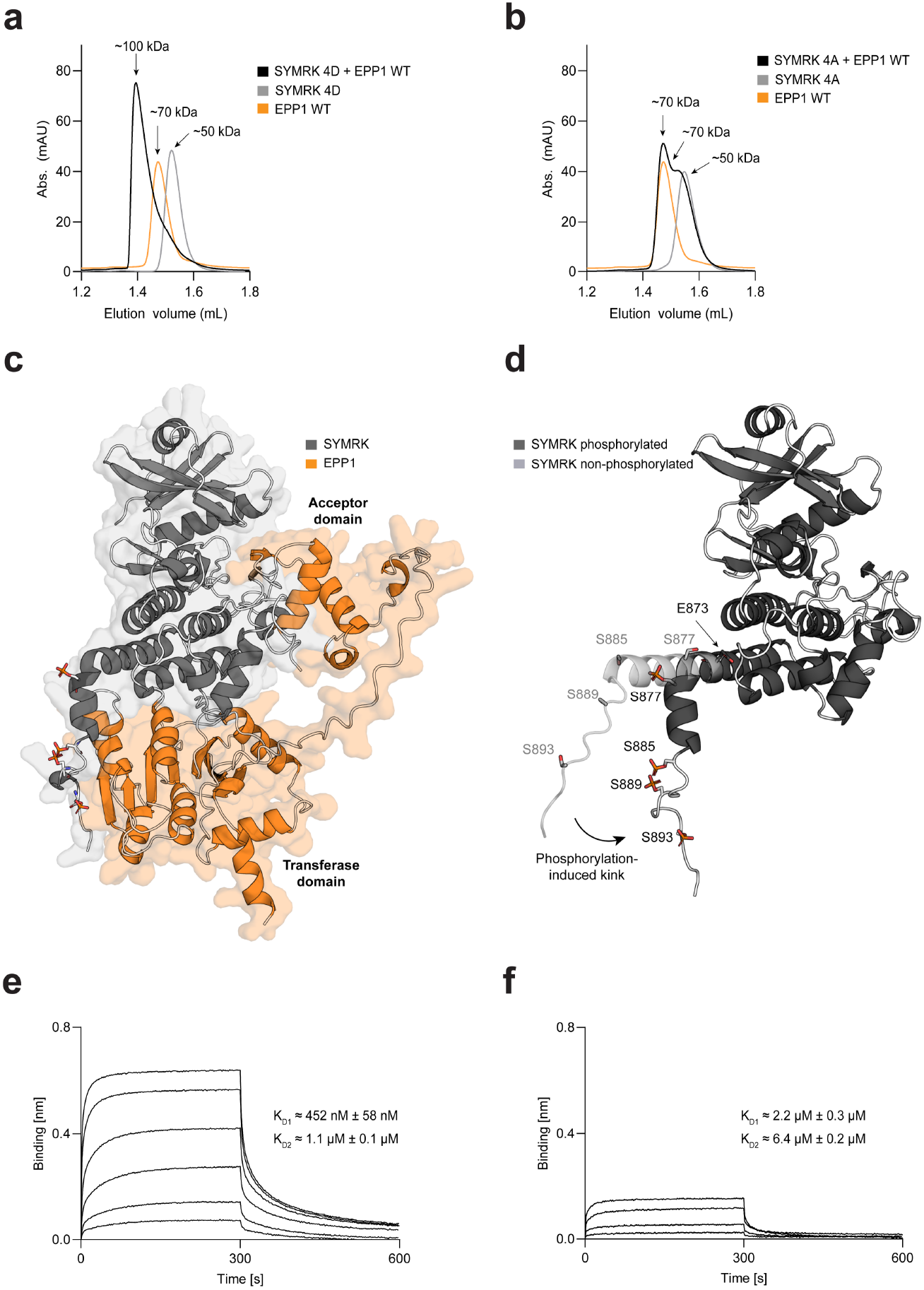
EPP1 recognizes the activated form of SYMRK. **a.** Analytic gel filtration profiles of SYMRK 4D (grey), EPP1a WT (orange) and a 1:1 mix of SYMRK 4D and EPP1 WT (black). **b**. Analytic gel filtration profiles of SYMRK 4A (grey), EPP1a WT (orange) and a 1:1 mix of SYMRK 4A and EPP1 WT (black). **c**. AlphaFold3 model of SYMRK (dark grey) phosphorylated on S877, S885, S889 and S893 forming a complex with *Lotus* EPP1a (orange). **d**. SYMRK modelled with phosphorylated residues (dark grey) or without (light grey). Phosphorylation of residue S877 in the alpha-I of SYMRK introduces a kink in the helix, likely due to electrostatic repulsion from residue E873. The resulting phosphorylation-induced kink creates a docking site for EPP1. **e**. Biolayer interferometry (BLI) sensorgram with immobilized SYMRK 4D and EPP1a WT concentration series. K_D_ values were calculated with a 2:1 heterogenous ligand fit. Representative traces, technical replicates and fits found in Extended Data Fig. 5. **f**. Biolayer interferometry (BLI) sensorgram with immobilized SYMRK 4A and EPP1a WT concentration series. K_D_ values were calculated with a 2:1 heterogenous ligand fit. Representative traces, technical replicates and fits found in Extended Data Fig. 6.

To gain structural insights into the observed complex between SYMRK 4D and EPP1, we initially attempted co-crystallization and single-particle cryo-electron microscopy, however, no structure was obtained. We therefore modelled the complex using AlphaFold3 which incorporates post-translational modifications such as phosphorylations in its structural predictions^18^. Activated SYMRK with phosphorylations at S877, S885, S889 and S893 forms a highly confident and tight complex with EPP1 (iPTM = 0.78) (Fig. 2c). SYMRK engages with EPP1 in two distinct interfaces: the acceptor domain of EPP1 contacts the N-lobe of SYMRK, whereas the transferase domain of EPP1 interacts with the phosphorylated C-lobe of SYMRK. This dual-site engagement is only observed in the phosphorylated and activated form of SYMRK, where phosphorylation of the alpha-I motif induces a kink in the extended SYMRK I-helix, creating a docking site for the EPP1 transferase domain (Fig. 2d). The acceptor domain of EPP1 simultaneously contacts the activation segment of SYMRK, placing EPP1 optimally for trans phosphorylation. Our data thus provide a structural framework for how activation of SYMRK allows the recruitment and formation of a stable complex with EPP1.

To quantify the interaction between activated SYMRK and EPP1, we performed biolayer interferometry (BLI). N-terminally Avi-tagged SYMRK variants (4D and 4A) were immobilized on Streptavidin-sensors and complex formation assessed using a concentration series of EPP1. Our experiments revealed distinct binding behaviors between the SYMRK variants and EPP1. For SYMRK 4D and EPP1, the sensorgrams displayed clear biphasic dissociation kinetics, reflecting binding of SYMRK 4D at two distinct interfaces of EPP1. The data were best fit using a 2:1 binding model, resulting in two dissociation constants; K_D1_ = 452 nM and K_D2_ = 1 μM (Fig. 2e, representative traces). In contrast, SYMRK 4A showed weaker binding to EPP1 with flatter binding curves, particularly at the lowest concentrations where complex formation could not be detected. The 2:1 model however still provided the best fit, yielding a K_D1_ = 2.2 μM and K_D2_ = 6.4 μM (Fig. 2f, representative traces). These results demonstrate that EPP1 exhibits a 5-6-fold higher binding affinity for activated SYMRK 4D compared to the non-activated SYMRK 4A. This affinity difference is critical for the formation of a stable and functional SYMRK-EPP1 signaling complex. Together, our data provide mechanistic understanding of how phosphorylation-dependent activation of SYMRK enables a high-affinity complex with EPP1, positioning EPP1 as a direct downstream signaling component of SYMRK in root nodule symbiosis.

### EPP1 is a direct substrate of SYMRK

To test whether SYMRK kinase activity is needed after EPP1 recruitment, we assessed the spontaneous nodulation phenotype of SYMRK 4D. A kinase-dead variant, SYMRK 4D K622E, failed to induce nodules (Fig. 3a), indicating that SYMRK kinase activity is required downstream of its own activation to propagate symbiotic signaling. To investigate whether EPP1 is a direct substrate of SYMRK, we focused on serine 77 (S77), previously identified as a phosphorylation site on EPP1^15–17^. We purified a phospho-ablated EPP1 S77A and performed *in vitro* trans-phosphorylation assays with different SYMRK versions. While EPP1 WT was phosphorylated by SYMRK and SYMRK 4A, phosphorylation by SYMRK 4D was markedly stronger, indicating that the phospho-mimetic 4D variant creates a high-affinity docking platform for EPP1 recruitment and efficient trans phosphorylation (Fig. 3b and Extended Data Fig. 7). In contrast, EPP1 S77A exhibited substantially reduced phosphorylation across all active SYMRK variants, indicating that EPP1 S77 is a primary target site of SYMRK transphosphorylation. Notably, even in the EPP1 S77A mutant, weak residual phosphorylation is observed -most prominently with SYMRK 4D -reflecting other possible phosphorylation sites *in vitro*. These results demonstrate that SYMRK directly phosphorylates EPP1 on S77, and that robust and efficient phosphorylation requires activation of the SYMRK alpha-I docking interface.

**Figure 3.**
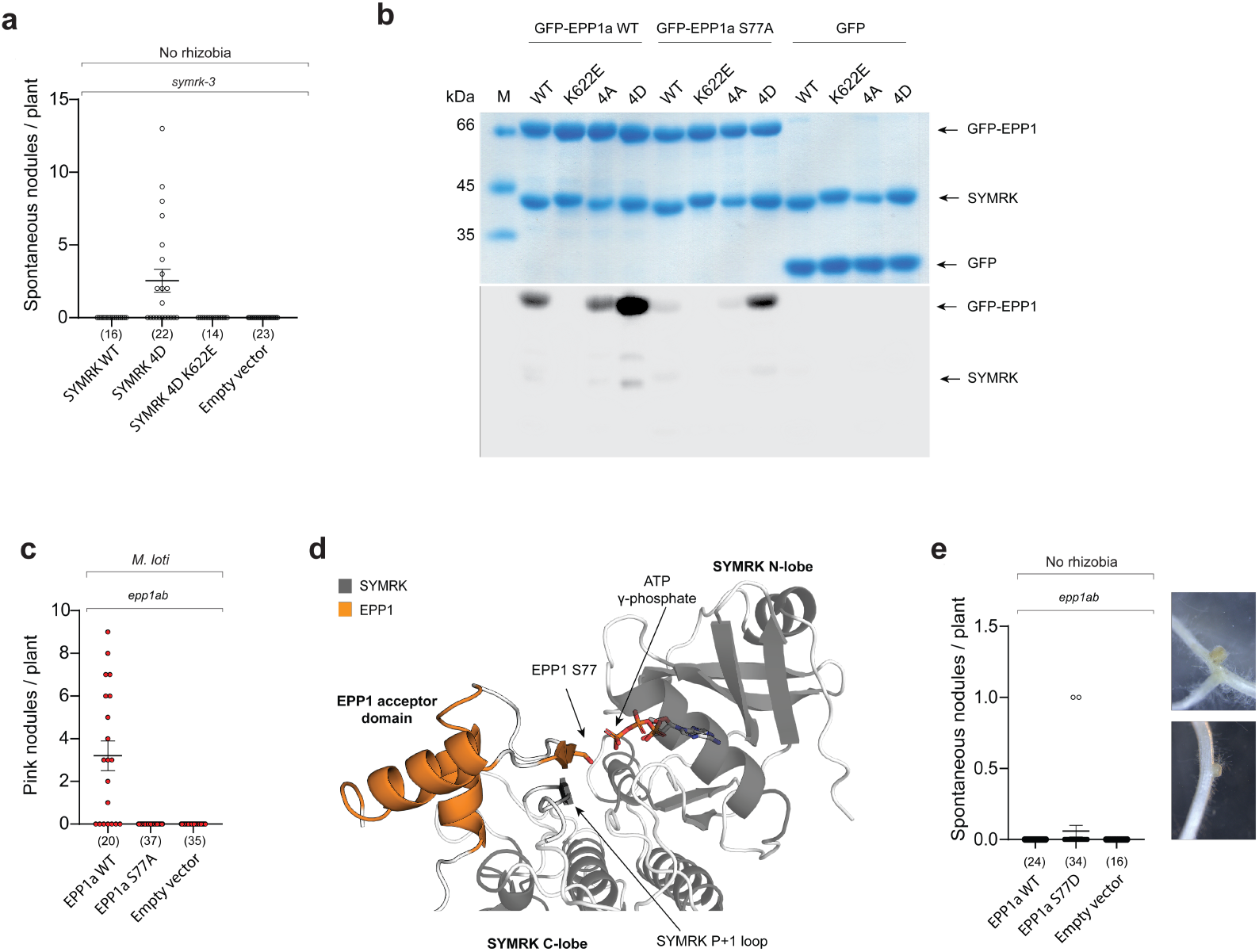
SYMRK phosphorylation of EPP1 on Serine 77 is critical for symbiosis. **a.** Number of white (uninfected) nodules on hairy roots expressing SYMRK WT, SYMRK 4D or SYMRK 4D K622E driven by its native promoter in *symrk-3* background 4 weeks after emerging of transgenic roots. The Numbers below graph specify the number of plants for each version of SYMRK or empty vector. **b**. Radioactive kinase activity assay with SYMRK variants and GFP-tagged EPP1a-variants. After incubation, the proteins were separated by SDS-PAGE (top) and radioactive γ-phosphate group transfer visualized by autoradiography (bottom). Free GFP was included as a negative control. **c**. Number of pink infected nodules 4 weeks after inoculation with *M. loti* rhizobia on *epp1a-1 epp1b-2* (*epp1ab*) mutants expressing indicated versions of EPP1 driven by its native promoter. The numbers below graph specify the number of plants for each version of EPP1 or empty vector. Note that the empty vector and SYMRK WT controls in **c** are the same data sets as in Fig. 1e. **d**. View of *Lotus* EPP1a WT (orange) presenting S77 of the acceptor domain to the P+1 loop of SYMRK (dark grey). SYMRK holds an ATP in its active site, displaying the γ-phosphate group ready for phospho-group transfer. **e**. Number of white (uninfected) nodules on hairy roots expressing EPP1a WT or EPP1a S77D driven by its native promoter in *epp1ab* background 4 weeks after emerging of transgenic roots. The numbers below graph specify the number of plants for each construct or empty vector.

To understand if phosphorylation of S77 is important for the function of EPP1, we next performed hairy root complementation experiments of the *epp1ab* double mutant with the phospho-ablated version of EPP1 S77A. EPP1 S77A is unable to rescue the loss of nodulation phenotype in *epp1ab* double mutant plants (Fig. 3c), showing that phosphorylation of S77 is essential for root nodule symbiosis signaling in *Lotus*. The identification of S77 in EPP1 as the primary phosphorylation site by SYMRK is mechanistically well-explained by the SYMRK-EPP1 structural model. Here, the EPP1 acceptor domain forms an anti-parallel β-sheet with the P+1 loop of SYMRK, positioning S77 ideally for trans-phosphorylation (Fig. 3d). This substrate alignment is a well-established mechanism for kinase-substrate interactions to ensure efficient and position-specific phospho-group transfer^20–22^.

Next, we investigated if an EPP1 S77D phospho-mimetic version could trigger downstream symbiotic signaling and tested this in hair root complementation experiments. Strikingly, EPP1 S77D driven by its native promoter produced spontaneous nodulation, albeit at low frequency (Fig. 3e), suggestion that phosphorylation of S77 is a key mechanism for signal propagation. Together, these results show that SYMRK autophosphorylation creates a docking site for EPP1, positioning its acceptor domain for optimal phosphorylation on S77 and that phosphorylation on S77 is essential and sufficient to trigger spontaneous nodulation in the absence of rhizobia.

### Recruitment of EPP1 by SYMRK connects the symbiotic signaling pathway

The structure of the SYMRK-EPP1 complex reveals how the phosphorylated, negatively charged alpha-I motif of SYMRK is recognized by the positively charged surface on the EPP1 transferase domain (Fig. 4a). This interaction explains the essential role of the alpha-I phosphorylation sites in symbiosis and suggests that recruitment of EPP1 is a key downstream consequence of SYMRK activation. To test whether EPP1 recruitment alone is sufficient to activate symbiotic signaling, we used a nanobody-based system to recruit EPP1 to SYMRK *in planta*. Strikingly, this synthetic recruitment triggered spontaneous nodule formation in the absence of rhizobia (Fig. 4b), showing that a nanobody-tethered SYMRK-EPP1 complex is sufficient to initiate nodule organogenesis and bypass the need for SYMRK activation. In contrast, nanobody-mediated recruitment of EPP1 to NFR1 did not induce nodulation (Fig. 4b), demonstrating that signaling specificity is tightly encoded in the pathway and that EPP1 functions specifically with SYMRK.

**Figure 4.**
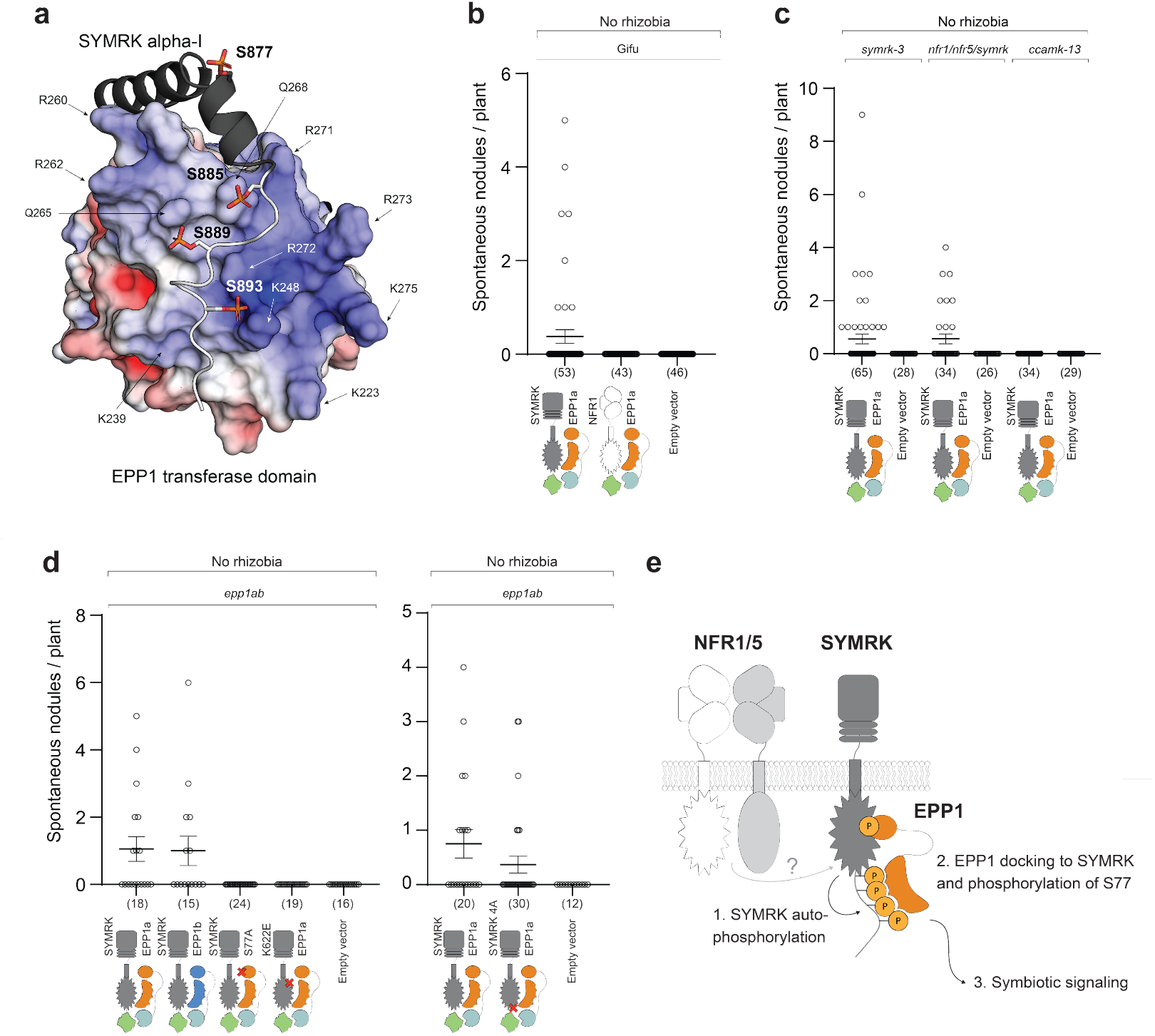
Engineered EPP1-SYMRK interaction drives symbiotic signaling. **a.** Visualization of the SYMRK alpha-I motif phosphorylated on S877, S885, S889 and S893 and its interaction with the EPP1 transferase domain. The EPP1 transferase domain has been visualized with surface charges, with blue indicating positively charged regions, grey indicating hydrophobic or uncharged regions, and red indicating negatively charged areas according to the electrostatic potential (±5 k_B_T/e). **b**. Number of white (uninfected) nodules on hairy roots expressing SYMRK or NFR1 synthetically linked to EPP1a via GFP-nanobody interaction 4 weeks after emerging of transgenic roots. All proteins are driven by their native promoter in Gifu WT background. The numbers below graph specify the number of plants for each construct or empty vector. **c**. Number of white (uninfected) nodules on hairy roots expressing SYMRK synthetically linked to EPP1a via GFP-nanobody interaction 4 weeks after emerging of transgenic roots. All proteins are driven by their native promoter in indicated mutant backgrounds. The numbers below graph specify the number of plants for each construct or empty vector. **d**. Number of white (uninfected) nodules on hairy roots expressing SYMRK WT, SYMRK 4A or kinase dead (K622E) artificially linked to EPP1a, EPP1b or EPP1 S77A via GFP-nanobody interaction 4 weeks after emerging of transgenic roots. All proteins are driven by their native promoter in *epp1ab* background. The numbers below graph specify the number of plants for each construct or empty vector. **e**. Suggested model for SYMRK activation and EPP1 phosphorylation. The potential activation of SYMRK by NFR1/5 leads to autophosphorylation of the alpha-I motif (step 1). EPP1 is recruited to activated SYMRK, leading to EPP1 phosphorylation on S77 (step 2) and progression of symbiotic signaling through CCaMK (step 3).

To define at which point EPP1 activation occurs within the symbiotic signaling pathway, we expressed nanobody-tethered SYMRK-EPP1 constructs in various symbiotic mutant backgrounds. In the *symrk* mutant, nanobody-tethered SYMRK-EPP1 drives spontaneous nodulation, indicating that downstream signaling can proceed in the absence of endogenous SYMRK. Strikingly, nodules also formed in the *nfr1/nfr5/symrk* triple mutant, demonstrating that once EPP1 is recruited to SYMRK, Nod factor receptors are no longer required for organogenesis signaling. This places EPP1 downstream of Nod factor perception. By contrast, no nodules formed in the *ccamk* mutant, revealing that the SYMRK-EPP1 signaling hub acts upstream of nuclear calcium decoding (Fig. 4c).

We next asked whether the requirement for SYMRK activation could be bypassed by nanobody-mediated recruitment of EPP1 to the inactivated SYMRK 4A (Fig. 4d). Nanobody-tethered SYMRK 4A-EPP1 still triggered spontaneous nodulation, indicating that the primary role of SYMRK alpha-I motif phosphorylation is to recruit EPP1 -a requirement that can be bypassed by directly associating the two proteins.

Finally, we investigated whether phosphorylation of EPP1 at S77 is still required when EPP1 is recruited to SYMRK via a nanobody. Interestingly, neither the phospho-ablated EPP1 S77A mutant nor the kinase-dead SYMRK K622E variant triggered nodule formation (Fig. 4d and Extended Data Fig. 8), indicating that recruitment alone is not sufficient and that EPP1 S77 phosphorylation must still occur to propagate symbiotic signal. These findings demonstrate that EPP1 functions downstream of SYMRK and requires its kinase activity to engage the symbiotic signaling pathway. Together, these results show that EPP1 operates within the common symbiotic signaling pathway and acts as a critical molecular link, connecting activated SYMRK to intracellular signal propagation (Fig. 4e). This places EPP1 as a central component in translating receptor activation at the plasma membrane into downstream responses in root nodule symbiosis.

## Discussion

In this study, we identify EPP1 as a crucial component in root nodule symbiosis, acting directly downstream of the activated SYMRK receptor. We demonstrate that phosphorylation of the SYMRK alpha-I motif creates a docking site that recruits EPP1 and positions the acceptor domain of EPP1 for optimal transphosphorylation by SYMRK. Our genetic and biochemical data show that SYMRK-dependent phosphorylation of EPP1 on S77 is essential for nitrogen-fixing symbiosis and sufficient to drive organogenesis signaling. Nanobody-mediated recruitment of EPP1 to SYMRK triggers spontaneous nodulation in the absence of rhizobia. These findings position EPP1 as a pivotal molecular decoder that translates the SYMRK phosphorylation status into activation of the intracellular symbiotic signaling cascade, thereby resolving a critical gap in our understanding of how receptor activation is coupled to downstream responses. Our study thus highlights EPP1 as a missing molecular link that connects perception of symbiotic signals at the plasma membrane to the activation of the cytoplasmic pathway.

## Methods

### Plant materials and growth conditions

Two independent LORE1 insertion lines were identified for *Epp1a* (LotjaGi1g1v0338600) and *Epp1b* (LotjaGi4g1v0221500) using the *LORE1* insertion mutant resource^23^. To generate double mutants, crosses were performed between *epp1a-1* and *epp1b-2*, as well as between *epp1a-2* and *epp1b-3*, respectively. Phenotyping analysis of the single and double *epp1* mutants was performed on agar plates supplemented with B&D nutrient solution^24^. Seedlings were inoculated with *M. loti* MAFF303099 expressing DsRED^25^ OD_600_ = 0.001 and grown 10 days for IT analysis and 4 weeks nodulation analysis. Hairy root experiments were carried out in *Lotus Japonicus* Gifu wild type, *epp1a-1 epp1b-2 double mutant (epp1ab), symrk-3, nfr1/nfr5/symrk*, or *ccamk-13* mutant backgrounds. Seeds were scarified with sandpaper here after surface sterilization for 15 min in 1% sodium hypochlorite. The seeds were set for germination on wet filter paper for 3 days. The seedlings were transferred to square agar plates with Gamborg’s B5 nutrient solution (Duchefa Biochemie). Hairy roots were performed on seedlings following^26^. Plants with transformed roots were transferred to magenta pots with LECA (Saint-Gobain Weber A/S) with B&D nutrient solution^24^. If indicated the plants were inoculated with *M. loti* MAFF303099 expressing DsRED^25^ OD_600_ = 0.01 and grown for 4 weeks. All plants were grown at 21 °C under a 16 h light/8 h dark cycle.

### Plasmid construction and cloning

For transgenic expression in *Lotus* roots, *Epp1a, Epp1b, Nfr1* and *SymRK* expression constructs were generated using synthesized modules of the genomic wild type and or mutated sequence of the respective genes and corresponding promoters (Thermo Fisher Scientific). For generating transgenic hairy roots, the pIV101 vector ^26^ was used. All plasmids were assembled using golden gate cloning as described in^27^. For transgenic expression in *Nicotiana benthamiana* leaves, EPP1a WT, SYMRK WT, SYMRK K622E, SYMRK 4A and SYMRK 4D were synthetized through Invitrogen GeneArt and codon-optimized for *Nicotiana benthamiana* expression. EPP1a and SYMRK variants were expressed under the control of the strong and constitutive *LjUbiquitin* promoter and tagged with 3xHA and SfGFP, respectively, in a binary vector using Golden Gate assembly. An overview of all modules used in this study are listed in Supplementary Table 1. All sequences for *E. coli* expression constructs used in this study are listed in Supplementary Table 2.

### Microscopy

Imaging of root hairs and ITs was performed using a Zeiss LSM 780 confocal microscope with a setup for bright field and DsRED, 561 nm excitation with 553 to 607 nm emission. Transgenic roots expressing indicated constructs were imaged using a Leica FluoStereo M165FC microscope equipped with a Leica DFC310 FX camera.

### Transient expression

Proteins were produced in leaves of 4-week-old *Nicotiana benthamiana* plants by *Agrobacterium*-mediated transformation using the AGL1 strain. Bacteria were grown in liquid LB media, pelleted (9,000 g, 3 min) and washed twice with infiltration buffer (10 mM MES-KOH pH 5.6, 10 mM MgCl_2_, 150 µM acetosyringone). Optical density at 600 nm (OD_600_) was adjusted to 0.25 for each strain, and the P19 silencing suppressor (OD_600_ = 0.2) was added to the mixture. Leaves were infiltrated with Agrobacterium using a needleless syringe, and samples were harvested 3 days post-infiltration.

### Co-immunoprecipitation

Leaf discs from *Nicotiana benthamiana* were frozen in liquid nitrogen and ground. Proteins were extracted at a 1:4 ratio (w:v) in extraction buffer (25 mM HEPES pH 7.5, 150 mM NaCl, 10 mM EDTA, 10% Glycerol, 1 mM DTT, 0.5% n-dodecyl-β-D-maltoside (DDM) supplemented with a protease inhibitor cocktail (Sigma)). Samples were solubilized for 30 min at 4°C and centrifuged (100,000g, 30 min, 4°C). The supernatant was incubated with 5 µL of magnetic agarose GFP-trap beads (ChromoTek) previously blocked for 1h with 1% BSA. Beads were washed three times with washing buffer (extraction buffer containing 0.25% DDM) and bound proteins were eluted by boiling in Laemmli at 95°C for 5 min. Western blotting was performed as previously described^28^.

### Structural modeling

Structural predictions were generated using the public AlphaFold3 server^18^ (v3.0.1, Deepmind/Isomorphic Labs, https://alphafoldserver.com/) with default parameters. For complex modeling, *Lotus* SYMRK residues 578-897 (phosphorylated on S877, S885, S889 and S893) and *Lotus* EPP1a residues 54-364 were modelled together. The top scoring complex yielded an interchain predicted TM-score (iPTM) of 0.78. For individual predictions, SYMRK residues 545-923 (with or without S877, S885, S889 and S893 phosphorylations) and EPP1a residues 1-364 were modelled. For all predictions, top-ranking models based on average pLDDT scores were selected for further analysis.

### Protein purification

Constructs for transgenic *E. coli* expression of SYMRK and EPP1 were synthesized and cloned into the pAH10R7Sumo3C^29^ and pET21b(+) vectors respectively by GenScript. All protein constructs were expressed in the Rosetta 2 *E. coli* strain (New England Biolabs) under identical conditions. Large scale cultures were grown in LB at 37 °C 110 rpm until OD_600_ = 0.8. Cultures were cold shocked followed by induction of protein expression with 0.4 mM IPTG at 18 °C 110 rpm overnight. Soluble his-tagged protein was initially purified from E. coli lysate on a Protino Ni-NTA column (Macherey-Nagel) equilibrated in buffer A (500 mM NaCl, 25 mM HEPES pH 7.5, 20 mM imidazole, 5 mM β-mercaptoethanol, 5% glycerol) and eluted with buffer B (500 mM NaCl, 25 mM HEPES pH 7.5, 500 mM imidazole, 5 mM β-mercaptoethanol, 5% glycerol). Eluates were dialyzed at 4 °C overnight (200 mM NaCl, 25 mM HEPES pH 7.5, 20 mM imidazole, 5 mM β-mercaptoethanol, 5% glycerol, 1 mM MnCl_2_) cleaved for affinity tags and dephosphorylated with 1:50 molar ratio of 3C and 1:200 molar ratio of λ-phosphatase, respectively. The samples were separated from cleaved tags and enzymes through a Protino Ni-NTA column equilibrated in buffer A. Final purification steps were performed on either Superdex 200 increase 10/300 or Superdex 75 increase 10/300 columns (Cytiva) equilibrated in gel filtration buffer (200 mM NaCl, 25 mM HEPES pH 7.5, 20 mM imidazole, 5 mM β-mercaptoethanol). Purity at each step of the purification was assessed with relevant fractions through sodium dodecyl sulfate gel polyacrylamide gel electrophoresis (SDS-PAGE).

### Analytic gel filtration

All proteins in analytic gel filtration assays had final concentrations of 1.5 ug/uL. For complex evaluation, SYMRK and EPP1 were incubated in a 1:1 molar ratio for 1h on ice before being spun down at 16,000 g for 20 mins and subjected to analytical gel filtration. All experiments were performed on a Superdex 200 increase 3.2/300 (Cytiva).

### BLI

Experiments were performed on an Octet RED96 (Fortébio) at 15 °C and 1000 rpm. Streptavidin biosensors (Sartorius) were pre-equilibrated for 20 mins in assay buffer (200 mM NaCl, 25 mM HEPES pH 7.5, 5 mM β-mercaptoethanol, 0.01% v/v Tween20). N-terminally Avi-tagged SYMRK 4D or 4A variants (15 ug/mL) were immobilized until a 1.5 nm response was reached, followed by a 120s washing step in assay buffer. Kinetics were measured over 300s association and 300s dissociation steps with a twofold dilution series of EPP1a WT (6 uM to 0.1875 uM). Unloaded sensors and sensors exposed only to buffer served as negative controls. Non-specific binding was subtracted and data aligned to average of pre-association baseline. Sensors were regenerated between each triplicate in 15s regeneration buffer (assay buffer + 1M NaCl), followed by 15s in assay buffer repeated three times. Kinetic parameters were determined using a 2:1 heterogenous ligand model in Octet Data Analysis HT 12.0 (Fortébio), with global fitting across all analyte concentrations within each replicate (grouped by sample ID and cycle). K_D_ values are reported as the mean ± SD across replicates. Raw sensorgrams were visualized in GraphPad Prism (version 10).

### Kinase assays

The assay was performed with all proteins in gel filtration buffer. Each reaction contained 2.5 ug of each protein and was initiated with the addition of 5 mM MgCl_2_ and 100 nCi γ-^32^P-labeled ATP. Reactions were left for 1h at room temperature and stopped by addition of 3 uL SDS loading dye and boiling at 95 °C for 5 mins. Samples were separated by SDS-PAGE, and the SDS-PAGE gel was subsequently allowed to incubate in an Autoradiography Hypercassette (Cytiva Amersham) overnight. The radiograph was developed on a Sapphire FL biomolecular imager (Azure Biosciences).

## Author contributions

M.M.M.N., N.B.A., J.S. and K.R.A. conceived the project. M.M.M.N., N.B.A., E.B. and D.L. performed experiments. M.M.M.N., N.B.A., E.B., D.L., J.S. and K.R.A. analyzed data. M.M.M.N., N.B.A. and K.R.A. wrote the manuscript with input from all authors.

## Acknowledgements

We would like to thank Majken K. Sørensen for plant care and seed production in our greenhouse, Maria Vinther for help with *E. coli* work, Emma Madsen for help with plant work and Mille Sonneby for help with protein work. This work was supported by the project Enabling Nutrient Symbioses in Agriculture (ENSA), that is funded by Bill & Melinda Gates Agricultural Innovations (INV-57461), the Bill & Melinda Gates Foundation and the Foreign, Commonwealth and Development Office (INV-55767), the Carlsberg Foundation grant (CF21-0139) and the Novo Nordisk Foundation (NNF24OC0094373, NNF24OC0095020, NNF24OC0095514).

