## Supplement for "EPP1 couples receptor activation to cytoplasmic signaling in root nodule symbiosis"

### Extended Data

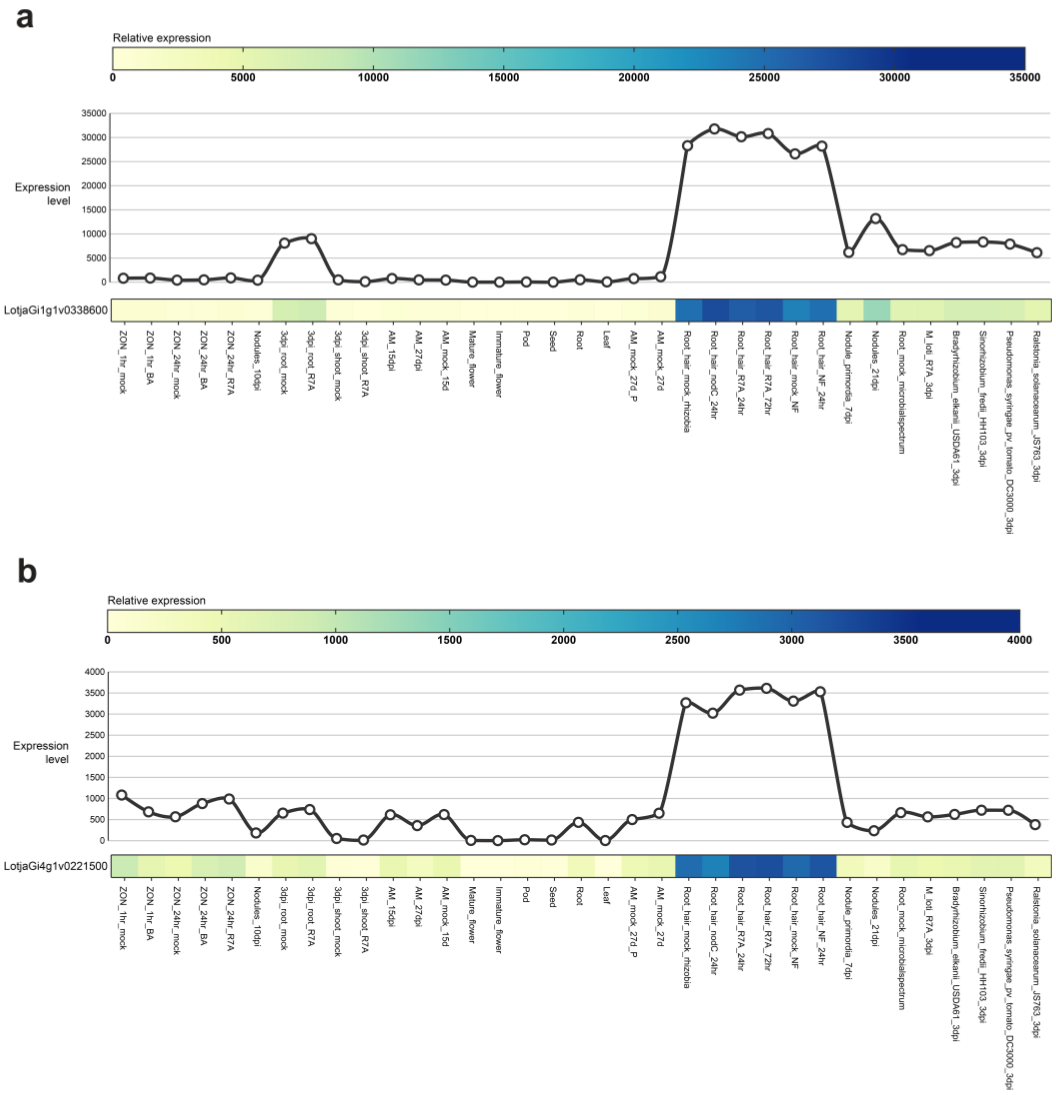

**Extended Data Fig. 1.** Lotus Base expression levels of Epp1a and Epp1b. **a**. Expression level of Epp1a (LotjaGi1g1v0338600.1). **b**. Expression level of Epp1b (LotjaGi4g1v0221500.1).

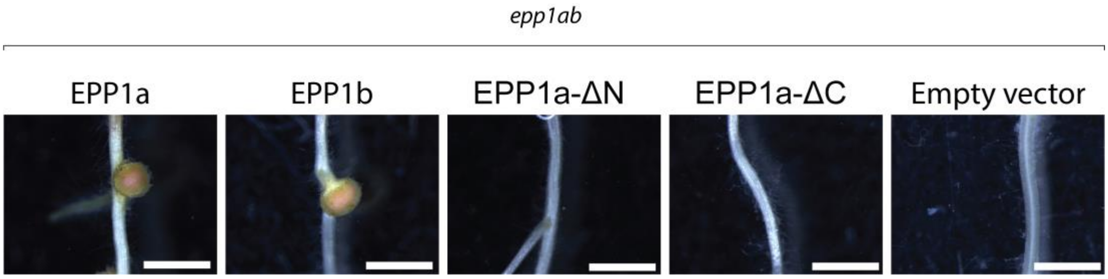

**Extended Data Fig. 2.** Representative pictures of nodules from **Fig. 1e**. Scalebar indicates 1 mm.

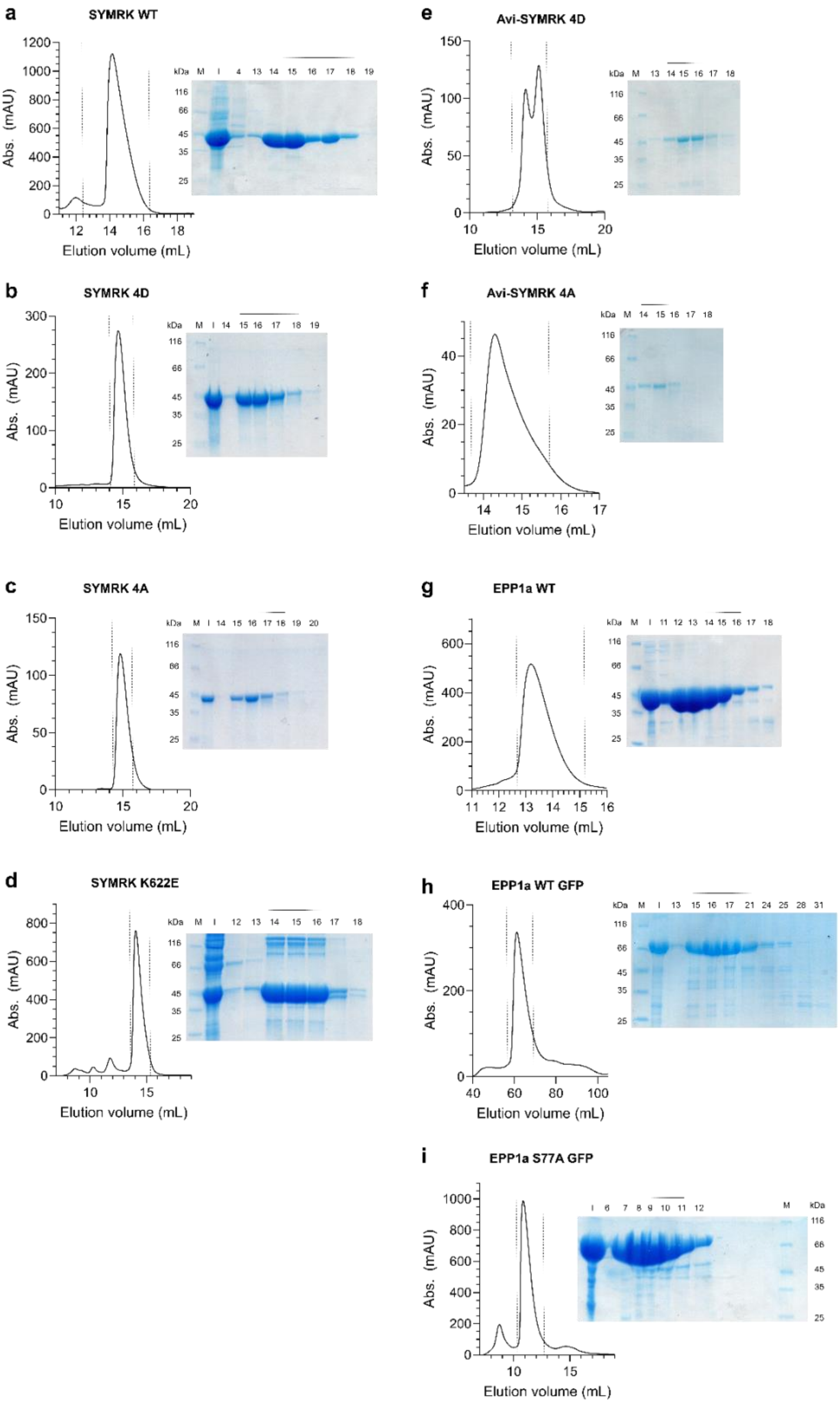

**Extended Data Fig. 3. (a-i)** Gel filtration profiles and SDS-PAGE gels for purifications of SYMRK and EPP1 constructs. Pooled fractions are indicated by dashed lines on chromatograms and a black bar over corresponding fractions in SDS-PAGE gels. M = marker, I = input.

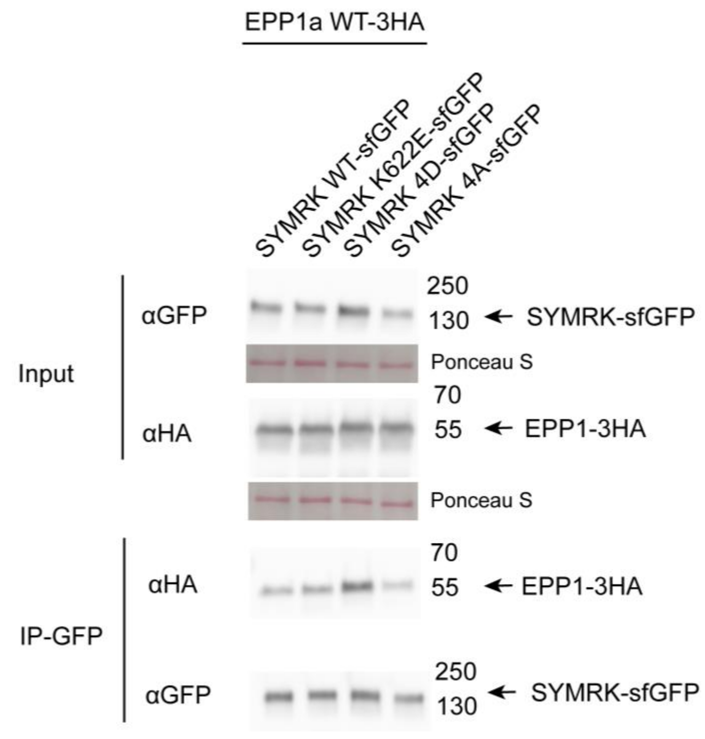

**Extended Data Fig. 4.** SYMRK-EPP1 Co-immunoprecipitation. The upper panel shows the input. *Lotus* SYMRK variants were immunoprecipitated using magnetic agarose GFP-trap beads, and the co-purified protein (*Lotus* EPP1a-3xHA) was detected using an anti-HA antibody (bottom panel). Ponceau S staining was used as loading control.

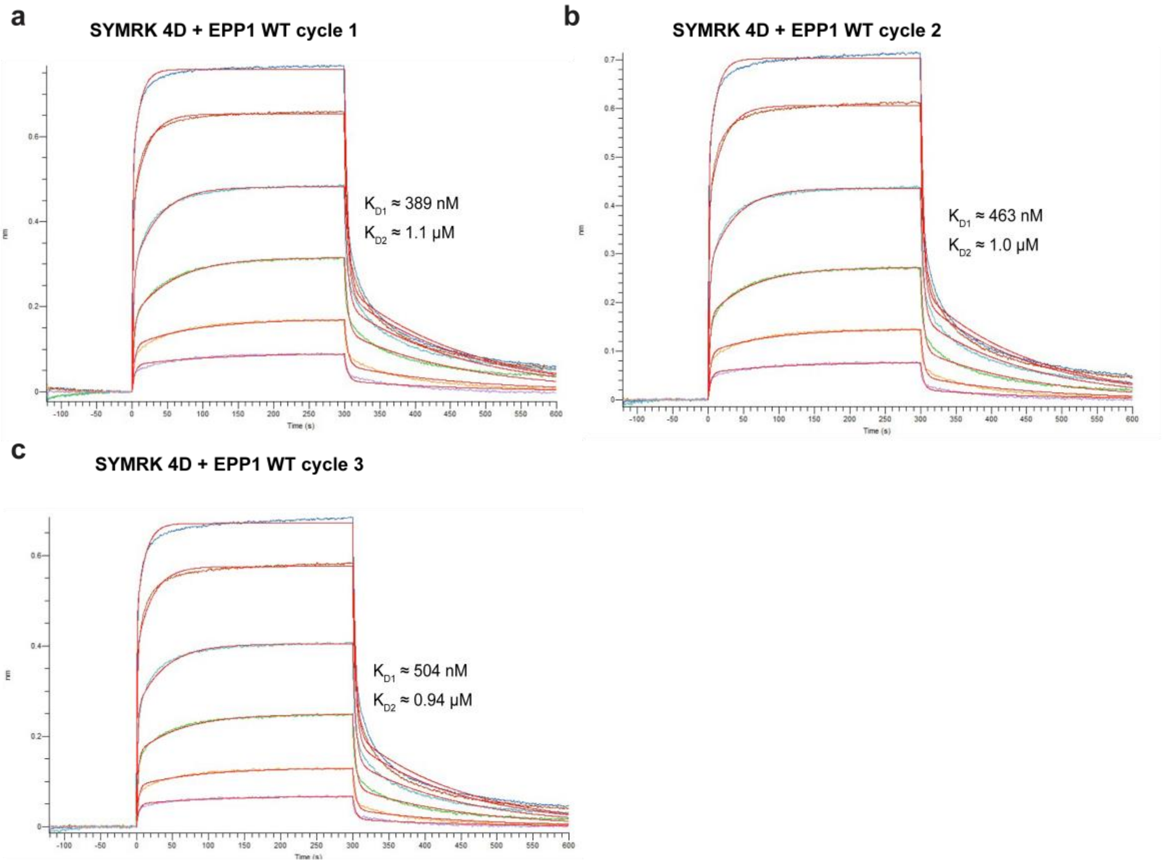

**Extended Data Fig. 5.** Biolayer interferometry replicates for SYMRK 4D - EPP1 WT binding. **a-c** Sensorgrams with experimental traces (coloured) and 2:1 heterogenous ligand fits (red) for replicate runs of SYMRK 4D and EPP1 WT in a two-fold concentration series (6 μM - 0.187 μM). **d**. Summary table of fitted kinetic parameters for each replicate.

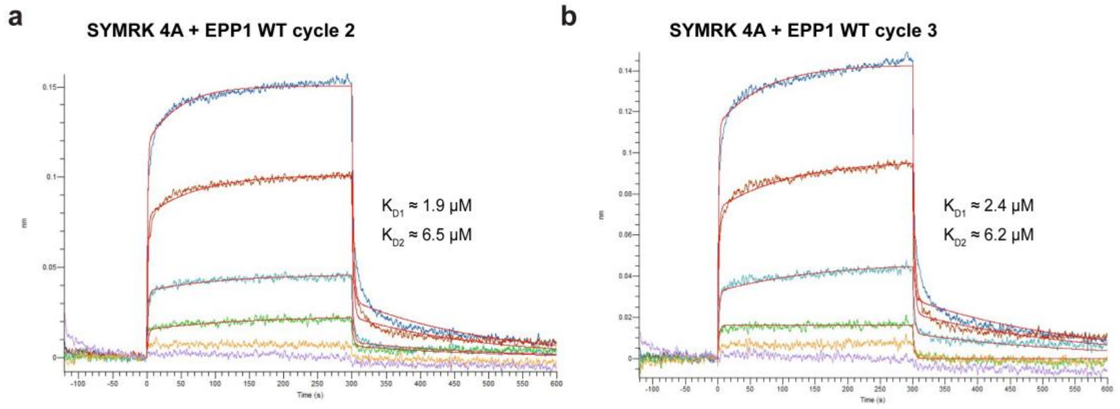

**Extended Data Fig. 6.** Biolayer interferometry replicates for SYMRK 4A - EPP1 WT binding. **a-b** Sensorgrams with experimental traces (coloured) and 2:1 heterogenous ligand fits (red) for replicate runs of SYMRK 4A and EPP1 WT in a two-fold concentration series (6 μM - 0.187 μM).

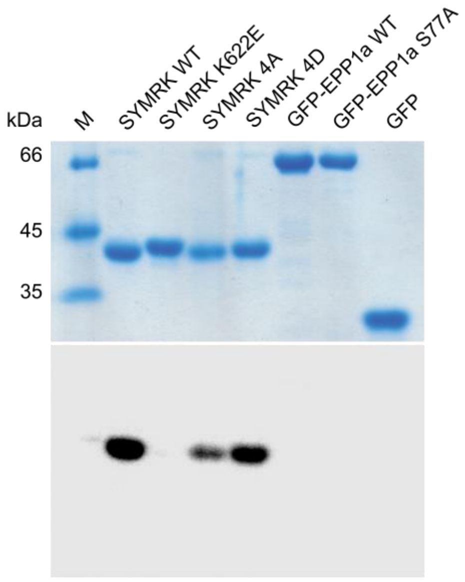

**Extended Data Fig. 7.** Radioactive kinase activity assay with SYMRK variants and GFP-tagged EPP1a-variants run separately. After incubation, the proteins were separated by SDS-PAGE (top) and radioactive γ-phosphate group transfer visualized by autoradiography (bottom). Free GFP was included as a negative control.

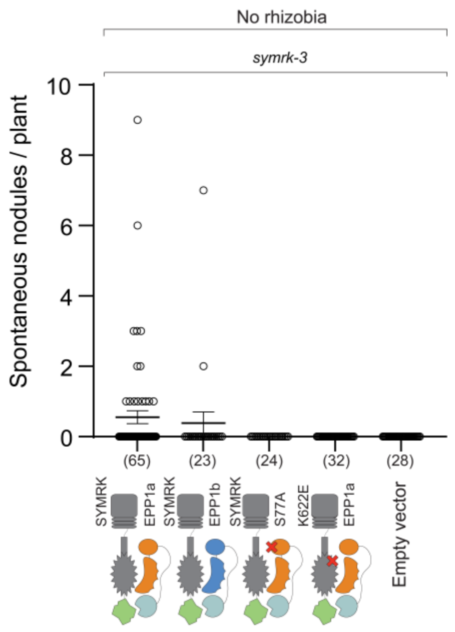

**Extended Data Fig. 8.** Number of white (uninfected) nodules on hairy roots expressing SYMRK WT or kinase dead (622E) synthetically linked to EPP1a, EPP1b or EPP177A via GFP-nanobody interaction 4 weeks after emerging of transgenic roots. All proteins are driven by their native promoter in *symrk-3* background. The numbers below graph specify the number of plants for each construct or empty vector. Note that the empty vector and SYMRK-EPP1a controls are the same data sets as Fig 4c.

### Supplementary information

| Construct name | Detailed name | Reference |
|---|---|---|
| **For generation of transgenic roots in *Lotus*** | | |
| WT EPP1a | *pEpp1a:Epp1a(WT)-t35S / pUBI:tYFP-NLS-t35s* | This study |
| WT EPP1b | *pEpp1b:Epp1b(WT)-t35S / pUBI:tYFP-NLS-t35s* | This study |
| EPP1a-ΔN | *pEpp1a:Epp1a(-ΔN)-t35S / pUBI:tYFP-NLS-t35s* | This study |
| EPP1a-ΔC | *pEpp1a:Epp1a(-ΔC)-t35S / pUBI:tYFP-NLS-t35s* | This study |
| EPP1a S77A | *pEpp1a:Epp1a(S77A)-t35S / pUBI:tYFP-NLS-t35s* | This study |
| EPP1a S77D | *pEpp1a:Epp1a(S77D)-t35S / pUBI:tYFP-NLS-t35s* | This study |
| WT SYMRK | *pSymRK:SymRK(WT)-mCherry-tSymRK / pUBI:tYFP-NLS-t35s* | *Abel et al 2024* |
| 4D SYMRK | *pSymRK:SymRK(4D)-mCherry-tSymRK / pUBI:tYFP-NLS-t35s* | *Abel et al 2024* |
| 4D SYMRK K622E | *pSymRK:SymRK(4D K622E)-mCherry-tSymRK / pUBI:tYFP-NLS-t35s* | This study |
| NBGFP-WT Epp1a / WT SYMRK-GFP | *pEpp1a:NBGFP-Epp1a(WT)-t35S / pSymRK:SymRK(WT)-GFP-tSymRK / pUBI:tYFP-NLS-t35s* | This study |
| NBGFP-WT Epp1a / WT NFR1-GFP | *pEpp1a:NBGFP-Epp1a(WT)-t35S / pNfr1:NFR1(WT)-GFP-tNfr1 / pUBI:tYFP-NLS-t35s* | This study |
| NBGFP-WT Epp1b / WT SYMRK-GFP | *pEpp1b:NBGFP-Epp1b(WT)-t35S / pSymRK:SymRK(WT)-GFP-tSymRK / pUBI:tYFP-NLS-t35s* | This study |
| NBGFP-S77A Epp1a / WT SYMRK-GFP | *pEpp1a:NBGFP-Epp1a(S77A)-t35S / pSymRK:SymRK(WT)-GFP-tSymRK / pUBI:tYFP-NLS-t35s* | This study |
| NBGFP-WT Epp1a / K622E SYMRK-GFP | *pEpp1a:NBGFP-Epp1a(WT)-t35S / pSymRK:SymRK(K622E)-GFP-tSymRK / pUBI:tYFP-NLS-t35s* | This study |
| **For *N. benthamiana* transformation** | | |
| WT EPP1a | *pUBI:Epp1a(WT)-3xHA-t35s* | This study |
| WT SYMRK | *pUBI:SymRK(WT)-sfGFP-t35s* | This study |
| K622E SYMRK | *pUBI:SymRK(K622E)-sfGFP-t35s* | This study |
| 4D SYMRK | *pUBI:SymRK(4D)-sfGFP-t35s* | This study |
| 4A SYMRK | *pUBI:SymRK(4A)-sfGFP-t35s* | This study |
| **For protein purification from E.coli** | | |
| WT EPP1a with/without N-term sfGFP | *pET21b(+)_10His_3C_sfGFP_TEV_EPP1a_WT_(res 1-364)* | This study |
| S77A EPP1a with/without N-term sfGFP | *pET21b(+)_10His_3C_sfGFP_TEV_EPP1a_S77A_(res 1-364)* | This study |
| WT SYMRK | *pAH10R7Sumo3C_SymRK_WT_(res 545-923)* | *Abel et al 2024* |
| 4D SYMRK | *pAH10R7Sumo3C_SymRK_4D_(res 545-923)_S877D_S885D_S889D_S893D* | *Abel et al 2024* |
| 4A SYMRK | *pAH10R7Sumo3C_SymRK_4A_(res 545-923)_S877A_S885A_S889A_S893A* | *Abel et al 2024* |
| K622E SYMRK | *pAH10R7Sumo3C_SymRK_K622E_(res 545-923)* | *Abel et al 2024* |
| Avi-SYMRK 4D | *pAH10R7Sumo3C_Avi_10xGS_SymRK_4D_(res 545-923)_S877D_S885D_S889D_S893D* | This study |
| Avi-SYMRK 4A | *pAH10R7Sumo3C_Avi_10xGS_SymRK_4A_(res 545-923)_S877A_S885A_S889A_S893A* | This study |

**Supplementary Table 1.** List of constructs used in this study.

***Lj*SYMRK WT (545-923)**
QKLIPWEGFAGKKYPMETNIIFSLPSKDDFFIKSVSIQAFTLEYIEVATERYKTLIGEGGFGSVYRGTLNDGQEVAVKVRSATSTQGT
REFDNELNLLSAIQHENLVPLLGYCNESDQQILVYPFMSNGSLQDRLYGEPAKRKILDWPTRLSIALGAARGLAYLHTFPGRSVIHR
DIKSSNILLDHSMCAKVADFGFSKYAPQEGDSYVSLEVRGTAGYLDPEYYKTQQLSEKSDVFSFGVVLLEIVSGREPLNIKRPRTE
WSLVEWATPYIRGSKVDEIVDPGIKGGYHAEAMWRVVEVALQCLEPFSTYRPSMVAIVRELEDALIIENNASEYMKSIDSLGGSNR
YSIVIEKRVLPSTTSTAESTITTQSLSHPQPR

***Lj*SYMRK 4D (545-923)**
QKLIPWEGFAGKKYPMETNIIFSLPSKDDFFIKSVSIQAFTLEYIEVATERYKTLIGEGGFGSVYRGTLNDGQEVAVKVRSATSTQGT
REFDNELNLLSAIQHENLVPLLGYCNESDQQILVYPFMSNGSLQDRLYGEPAKRKILDWPTRLSIALGAARGLAYLHTFPGRSVIHR
DIKSSNILLDHSMCAKVADFGFSKYAPQEGDSYVSLEVRGTAGYLDPEYYKTQQLSEKSDVFSFGVVLLEIVSGREPLNIKRPRTE
WSLVEWATPYIRGSKVDEIVDPGIKGGYHAEAMWRVVEVALQCLEPFSTYRPSMVAIVRELEDALIIENNADEYMKSIDDLGGDNR
YDIVIEKRVLPSTTSTAESTITTQSLSHPQPR

***Lj*SYMRK 4A (545-923)**
QKLIPWEGFAGKKYPMETNIIFSLPSKDDFFIKSVSIQAFTLEYIEVATERYKTLIGEGGFGSVYRGTLNDGQEVAVKVRSATSTQGT
REFDNELNLLSAIQHENLVPLLGYCNESDQQILVYPFMSNGSLQDRLYGEPAKRKILDWPTRLSIALGAARGLAYLHTFPGRSVIHR
DIKSSNILLDHSMCAKVADFGFSKYAPQEGDSYVSLEVRGTAGYLDPEYYKTQQLSEKSDVFSFGVVLLEIVSGREPLNIKRPRTE
WSLVEWATPYIRGSKVDEIVDPGIKGGYHAEAMWRVVEVALQCLEPFSTYRPSMVAIVRELEDALIIENNAAEYMKSIDALGGANR
YAIVIEKRVLPSTTSTAESTITTQSLSHPQPR

***Lj*SYMRK K622E (545-923)**
QKLIPWEGFAGKKYPMETNIIFSLPSKDDFFIKSVSIQAFTLEYIEVATERYKTLIGEGGFGSVYRGTLNDGQEVAVEVRSATSTQGT
REFDNELNLLSAIQHENLVPLLGYCNESDQQILVYPFMSNGSLQDRLYGEPAKRKILDWPTRLSIALGAARGLAYLHTFPGRSVIHR
DIKSSNILLDHSMCAKVADFGFSKYAPQEGDSYVSLEVRGTAGYLDPEYYKTQQLSEKSDVFSFGVVLLEIVSGREPLNIKRPRTE
WSLVEWATPYIRGSKVDEIVDPGIKGGYHAEAMWRVVEVALQCLEPFSTYRPSMVAIVRELEDALIIENNASEYMKSIDSLGGSNR
YSIVIEKRVLPSTTSTAESTITTQSLSHPQPR

**Avi-10xGS-*Lj*SYMRK 4D (545-923)**
GLNDIFEAQKIEWHESGGGSGGGSGQKLIPWEGFAGKKYPMETNIIFSLPSKDDFFIKSVSIQAFTLEYIEVATERYKTLIGEGGFG
SVYRGTLNDGQEVAVKVRSATSTQGTREFDNELNLLSAIQHENLVPLLGYCNESDQQILVYPFMSNGSLQDRLYGEPAKRKILDW
PTRLSIALGAARGLAYLHTFPGRSVIHRDIKSSNILLDHSMCAKVADFGFSKYAPQEGDSYVSLEVRGTAGYLDPEYYKTQQLSEK
SDVFSFGVVLLEIVSGREPLNIKRPRTEWSLVEWATPYIRGSKVDEIVDPGIKGGYHAEAMWRVVEVALQCLEPFSTYRPSMVAIV
RELEDALIIENNADEYMKSIDDLGGDNRYDIVIEKRVLPSTTSTAESTITTQSLSHPQPR

**Avi-10xGS-*Lj*SYMRK 4A (545-923)**
GLNDIFEAQKIEWHESGGGSGGGSGQKLIPWEGFAGKKYPMETNIIFSLPSKDDFFIKSVSIQAFTLEYIEVATERYKTLIGEGGFG
SVYRGTLNDGQEVAVKVRSATSTQGTREFDNELNLLSAIQHENLVPLLGYCNESDQQILVYPFMSNGSLQDRLYGEPAKRKILDW
PTRLSIALGAARGLAYLHTFPGRSVIHRDIKSSNILLDHSMCAKVADFGFSKYAPQEGDSYVSLEVRGTAGYLDPEYYKTQQLSEK
SDVFSFGVVLLEIVSGREPLNIKRPRTEWSLVEWATPYIRGSKVDEIVDPGIKGGYHAEAMWRVVEVALQCLEPFSTYRPSMVAIV
RELEDALIIENNAAEYMKSIDALGGANRYAIVIEKRVLPSTTSTAESTITTQSLSHPQPR

***Lj*EPP1a (1-364)**
MGTNKQEHEISDPGSEVTNQEKVENEVKDTPKLATHKIITESNKSNSMVIKKRHTLIPPHIIAEAISSIRDIDIRWSGPITPKEMEYVE
QYVLAKYPEYEGLIEGDGNGVDMSMFMINEEPSEYDRGKSPCGTPSPRESSAYLFGTNLPETDRTKIQLEPSRLLDILNKKSSFP
GSFISIPEIQVRNKVLKHYGLPDEEYLVLFTPSYKDAMMLVGESYPFLKGNFYMTILDQEEDYIKEFASFKESKVILAPKTWLDLRIR
GSQLSQNFRRRCKISSKGLFSYPSDASGTMHWISEAHRNSWHVLLDASAFVVGKDRLHLALHRPDFVVCSLDNTHSNPSRITCL
LVRKKSFDTSGASSQVVE

**10xHis-3C-GFP-TEV-*Lj*EPP1a (1-364)**
MGSHHHHHHHHHHGSLEVLFQGPGSGSKGEELFTGVVPILVELDGDVNGHKFSVRGEGEGDATNGKLTLKFICTTGKLPVPWP
TLVTTLTYGVQCFARYPDHMKQHDFFKSAMPEGYVQERTISFKDDGTYKTRAEVKFEGDTLVNRIELKGIDFKEDGNILGHKLEYN
FNSHNVYITADKQKNGIKANFKIRHNVEDGSVQLADHYQQNTPIGDGPVLLPDNHYLSTQSVLSKDPNEKRDHMVLLEFVTAAGIT
HGMDELYKSGSGSENLYFQGGGSGSMGTNKQEHEISDPGSEVTNQEKVENEVKDTPKLATHKIITESNKSNSMVIKKRHTLIPPHII
AEAISSIRDIDIRWSGPITPKEMEYVEQYVLAKYPEYEGLIEGDGNGVDMSMFMINEEPSEYDRGKSPCGTPSPRESSAYLFGTNL
PETDRTKIQLEPSRLLDILNKKSSFPGSFISIPEIQVRNKVLKHYGLPDEEYLVLFTPSYKDAMMLVGESYPFLKGNFYMTILDQEED
YIKEFASFKESKVILAPKTWLDLRIRGSQLSQNFRRRCKISSKGLFSYPSDASGTMHWISEAHRNSWHVLLDASAFVVGKDRLHL
ALHRPDFVVCSLDNTHSNPSRITCLLVRKKSFDTSGASSQVVE

**10xHis-3C-GFP-TEV-*Lj*EPP1a S77A (1-364)**
MGSHHHHHHHHHHGSLEVLFQGPGSGSKGEELFTGVVPILVELDGDVNGHKFSVRGEGEGDATNGKLTLKFICTTGKLPVPWP
TLVTTLTYGVQCFARYPDHMKQHDFFKSAMPEGYVQERTISFKDDGTYKTRAEVKFEGDTLVNRIELKGIDFKEDGNILGHKLEYN
FNSHNVYITADKQKNGIKANFKIRHNVEDGSVQLADHYQQNTPIGDGPVLLPDNHYLSTQSVLSKDPNEKRDHMVLLEFVTAAGIT
HGMDELYKSGSGSENLYFQGGGSGSMGTNKQEHEISDPGSEVTNQEKVENEVKDTPKLATHKIITESNKSNSMVIKKRHTLIPPHII
AEAISSIRDIDIRWAGPITPKEMEYVEQYVLAKYPEYEGLIEGDGNGVDMSMFMINEEPSEYDRGKSPCGTPSPRESSAYLFGTNL
PETDRTKIQLEPSRLLDILNKKSSFPGSFISIPEIQVRNKVLKHYGLPDEEYLVLFTPSYKDAMMLVGESYPFLKGNFYMTILDQEED
YIKEFASFKESKVILAPKTWLDLRIRGSQLSQNFRRRCKISSKGLFSYPSDASGTMHWISEAHRNSWHVLLDASAFVVGKDRLHL
ALHRPDFVVCSLDNTHSNPSRITCLLVRKKSFDTSGASSQVVE

**Supplementary Table 2.** Sequences of *E. coli* expression constructs used in this study.
